# Incentive and Dopamine Sensitization Produced by Intermittent but Not Long Access Cocaine Self-Administration

**DOI:** 10.1101/499475

**Authors:** Alex B. Kawa, Alec C. Valenta, Robert T. Kennedy, Terry E. Robinson

## Abstract

Recent studies suggest that the temporal pattern of drug use (pharmacokinetics) has a profound effect on the ability of self-administered cocaine to produce addiction-like behavior in rodents, and to change the brain. To further address this issue, we compared the effects of Long Access (LgA) cocaine self-administration, which is widely used to model the transition to addiction, with Intermittent Access (IntA), which is thought to better reflect the pattern of drug use in humans, on the ability of self-administered cocaine to increase dopamine (DA) overflow in the core of the nucleus accumbens (using in vivo microdialysis), and to produce addiction-like behavior. IntA experience was more effective than LgA in producing addiction-like behavior – a drug experience-dependent increase in motivation for cocaine assessed using behavioral economic procedures, and cue-induced reinstatement – despite much less total drug consumption. There were no group differences in basal levels of DA in dialysate, but a single self-administered IV injection of cocaine increased DA in the core of the nucleus accumbens to a greater extent in rats with prior IntA experience than those with LgA or Short Access (ShA) experience, and the latter two groups did not differ. Furthermore, high motivation for cocaine was associated with a high DA response. Thus, IntA, but not LgA, produced both incentive and DA sensitization. This is consistent with the notion that a hyper-responsive dopaminergic system may contribute to the transition from casual patterns of drug use to the problematic patterns that define addiction.

## INTRODUCTION

Drug self-administration procedures have proved invaluable for isolating how drug use can change the brain in ways that promote the development of addiction (Weeks, 1962; Ahmed, 2018). However, mere drug-taking does not necessarily produce the loss of control and aberrant motivation that defines addiction. For this reason there has been considerable interest in the development of self-administration procedures and conditions (here we focus on cocaine) that most reliably produce addiction-like behavior (Ahmed & Koob, 1998; Deroche-Gamonet *et al*., 2004; Vanderschuren & Everitt, 2004; Ahmed, 2012; Zimmer *et al*., 2012). To this end, Ahmed and Koob (1998) first introduced what has become known as the Long Access (LgA) self-administration procedure, which is arguably the most widely used rodent model of addiction. During LgA, rats are typically allowed to self-administer cocaine on a Fixed Ratio schedule for 6+ hours/daily session; and relative to rats allowed to self-administer cocaine for only 1-2 hours/session (Short Access, ShA), LgA experience is reported to result in the emergence of a number of addiction-like behaviors, including escalation of intake, increased motivation for drug, and continued drug-seeking in the face of an adverse consequence (for reviews see, Ahmed, 2012; Edwards & Koob, 2013).

Studies on the neurobiological consequences of LgA cocaine experience suggest that LgA decreases dopamine (DA) neurotransmission in the nucleus accumbens core (relative to ShA). For example, LgA experience decreases cocaine-induced inhibition of DA uptake, electrically evoked DA release when measured *ex vivo,* and cocaine-induced DA overflow *in vivo* measured with microdialysis (Ferris *et al*., 2011; Calipari *et al*., 2013, 2014). In addition, increasing LgA cocaine self-administration experience has been associated with a progressive decrease in DA release accompanying operant responding measured using voltammetry *in vivo* (Willuhn *et al*., 2014). Although there are exceptions (e.g., Ahmed *et al*., 2003), these findings have been interpreted as support for the idea that cocaine addiction is due, in part, to a cocaine-induced *hypo*-dopaminergic state, and continued drug-seeking is motivated by a desire to overcome this DA deficiency (e.g., Caprioli *et al*., 2014; Koob & Volkow, 2016; U.S. Department of Health and Human Services, 2016; Volkow *et al*., 2016).

A fundamental assumption of the LgA model is that the amount of drug exposure is the critical factor in the emergence of addiction-like behavior. As stated by Ahmed (2012), “*addiction-causing neuropathological processes could be set in motion only when rats can expose themselves sufficiently to cocaine to cross the ‘threshold of addiction*’ … *below this critical level of cocaine exposure, there would be no drug-induced neuropathological changes, and drug use would remain under control, at least in the majority of drug-exposed individuals*” (p. 110). However, studies using a more recently introduced model of the transition to cocaine addiction have challenged this assumption (Zimmer et al. 2011, 2012; Allain et al. 2015).

During a LgA session rats first ‘load up’ and then maintain relatively high brain concentrations of cocaine for the duration of the 6+ hour session. However, Zimmer et al. (2012) noted that this may not reflect the pattern of cocaine use in humans, which is much more intermittent both between and *within* bouts of use. Thus, Zimmer et al. (2012) developed what they called the Intermittent Access (IntA) cocaine self-administration procedure to better model the repeated spikes in brain cocaine concentrations thought to reflect the pattern of use in humans within a bout of use, especially during the transition to addiction. To this end, rats were allowed 5 min of unlimited access to cocaine followed by a 25 min period when drug was not available, and this repeated throughout the session, resulting in successive spikes in brain cocaine concentrations. The important finding was that IntA experience produced greater motivation for cocaine than LgA, despite much less total drug consumption. There are now a number of reports that IntA not only increases motivation for cocaine to a greater extent than LgA, but is more effective in producing a number of other addiction-like behaviors (Kawa *et al*., 2016; Allain *et al*., 2017, 2018; Allain & Samaha, 2018; James *et al*., 2018; Kawa & Robinson, 2018).

Furthermore, unlike LgA, which blunts DA neurotransmission, several studies have suggested that IntA does exactly the opposite; it sensitizes DA neurotransmission. For example, in brain slices IntA is reported to increase the ability of cocaine to inhibit DA uptake and to increase electrically-evoked DA release (Calipari *et al*., 2013). Of course, this raises questions concerning the role of a hypo- vs *hyper*-dopaminergic state in addiction, especially given that IntA is more effective than LgA in producing addiction-like behavior. However, to date, the only studies on the effects of IntA on DA neurotransmission have been in brain slices, *ex vivo*. Therefore, given the theoretical implications, it is important to determine whether IntA cocaine experience produces similar effects in freely moving rats, *in vivo*. That was the purpose of the present experiment, in which we used *in vivo* microdialysis in freely moving rats to quantify changes in DA (and other analytes) in response to a single self-administered injection of cocaine, following ShA, LgA or IntA cocaine self-administration experience.

## MATERIALS AND METHODS

A total of 50 male Sprague-Dawley rats (Envigo, Haslett, MI) weighing 250-275 g upon arrival were housed individually in a climate-controlled colony room, on a reverse 12-h light/12-h dark cycle (lights on at 20:00). All testing was conducted during the 12-hour lights off period. After arrival, rats were given 1 week to acclimate to the colony room before surgery. Water and food were available *ad libitum* until 2 days before the first day of self-administration, at which point the rats were mildly food restricted to maintain a stable body weight for the remainder of the experiment (20-24 grams/day). There is evidence that maintaining adult, male rats at a stable body weight is more healthy than *ad libitum* feeding (Rowland, 2007). All procedures were approved by the University of Michigan Committee on the Use and Care of Animals (UCUCA).

### Apparatus

Behavioral testing was conducted in standard (22×18×13 cm) test chambers (Med Associates, St Albans, VT, USA) located inside sound-attenuating cabinets. A ventilating fan masked background noise. Within the test chambers, two nose poke ports were located 3 cm above the floor on the left and right side of the front wall, and one port was designated active and the other inactive (side counter-balanced across chambers). A red house light was located at the top, center of the back wall opposite the nose ports. During self-administration, intravenous cocaine infusions were delivered by a pump mounted outside the sound attenuating cabinet, through a tube connected to the rat’s catheter back port. The infusion tube was suspended into the chamber via a swivel mechanism, allowing the rat free movement. All measures were recorded using Med Associates software.

Microdialysis test sessions were conducted in separate but identical chambers to those described above. The only difference was that the swivel system in these boxes allowed for the microdialysis inlet tubing, outlet tubing, and the drug delivery tubing to be connected simultaneously.

### Intravenous catheter surgery

Rats underwent intravenous catheter surgery as described previously (Crombag *et al*., 2000). Briefly, rats were anesthetized using ketamine hydrochloride (90 mg/kg i.p.) and xylazine (10mg/kg i.p.) and a catheter was inserted into the right jugular vein and tubing was run subcutaneously to a port located on the rat’s back. During recovery from surgery rats were administered the analgesic carprofen (5 mg/kg s.c.). Following surgery, catheters were flushed daily with 0.2 ml sterile saline containing 5 mg/ml gentamicin sulfate (Vedco, MO). Catheter patency was tested periodically with intravenous injection of 0.1 ml methohexital sodium (10 mg/ml in sterile water, JHP Pharmaceuticals). If a rat did not become ataxic within 10 seconds of the injection, the catheter was considered not patent and the rat was removed from the study.

### Self-administration: acquisition

Rats were given ~7 days to recover from the catheter surgery, and then self-administration training commenced (Fig. 1a). The rats were placed into the chamber with the house light illuminated. After 2 minutes the house light was extinguished and this signaled the beginning of each session. At that time a nose poke into the active port, detected by an infrared photo beam, resulted in an intravenous infusion of cocaine hydrochloride (NIDA) dissolved in 0.9% sterile saline (0.4 mg/kg/infusion, weight of the salt, in 50 μl delivered over 2.6 seconds) on a Fixed Ratio-1 (FR-1) schedule. Each infusion was paired with the illumination of a cue light in the active nose port for 20 seconds. Nose pokes during this 20 sec period were recorded but had no consequences. An inactive nose port was also present at all times and pokes there were recorded but had no consequences. To ensure that during initial training all rats received the same amount of cocaine and cue exposure an infusion criteria (IC) procedure was used, as described previously (Saunders & Robinson, 2010). During these acquisition sessions, session length was determined by how long it took each rat to reach the predetermined number of infusions, not by an explicit time limit. Each rat had 2 sessions at IC10 (10 infusions), 3 sessions at IC20, and 4 sessions at IC40. A total of 5 rats were excluded during acquisition training because they failed to reach the infusion criteria or failed to discriminate between the active nose port and the inactive nose port.

**Fig. 1.**
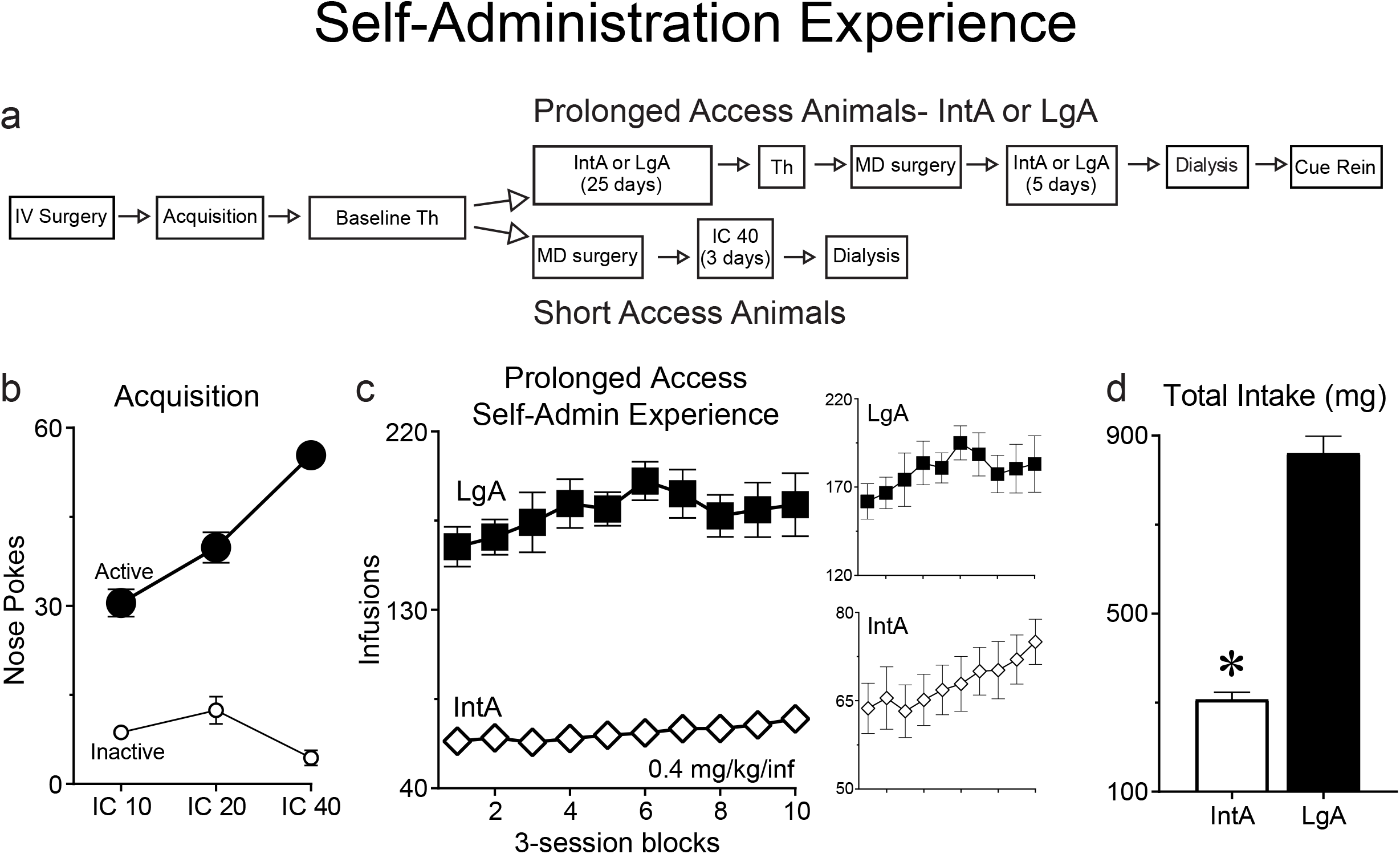
The flow diagram shows the overall experimental design and timeline for the experiment (**a**) (see methods for details). An Infusion Criterion (**IC**) procedure was used for the acquisition of cocaine self-administration (**b**). IntA rats and LgA rats both escalated their intake/session to a similar extent across all 30 self-administration sessions (**c**). LgA rats consumed far more cocaine on average than IntA rats over the course of 30 self-administration sessions under their respective procedures (**d**). (Th: Threshold Test, IntA: Intermittent Access, LgA: Long Access, IV/MD surgery: Intravenous catheter/Microdialysis guide cannula surgery). In panel c each point represents the average of three consecutive self-administration sessions. See methods for the number of rats included in each group and panel. Values represent means ± SEMs.

### Self-administration: within-session threshold procedure

The day after the final acquisition session, all rats were trained on a within-session threshold procedure and demand curves were generated, as described previously (Bentzley *et al*., 2013; Kawa *et al*., 2016). Briefly, each session (one per day) was 110 minutes in length, FR-1 throughout, and every 10 minutes the dose of drug was decreased on a quarter logarithmic scale (1.28, 0.72, 0.40, 0.23, 0.13, 0.072, 0.040, 0.023, 0.013, 0.007, 0.004 mg/kg/infusion). During the threshold procedure, the nose port cue light was illuminated for the duration of each infusion. Each rat was tested daily for at least four sessions and until it produced three consecutive sessions with less than 25% variation in α. For baseline data analysis α, P_Max_, and Q_0_ values (see below) were averaged over these last 3 sessions for each rat. A total of 3 rats were excluded for failure to stabilize or loss of catheter patency.

The threshold procedure yields a number of behavioral economic metrics. **Q_0_** is a theoretical measure of consumption when no effort is required; that is, an inherent extrapolation of the individual’s consumption at very low prices (Hursh & Silberberg, 2008; Oleson *et al*., 2011; Bentzley *et al*., 2013). **P_Max_** is defined as the price that elicits maximum responding; i.e., the maximum price (in effort) an individual is willing to pay to maintain Q_0_ (Hursh, 1991; Bentzley *et al*., 2013). Consumption remains relatively stable at prices lower than P_Max_ but falls rapidly at prices higher than P_Max_. Finally, **α** is a measure of demand elasticity and is equivalent to the slope of the demand curve (Hursh & Silberberg, 2008; Bentzley *et al*., 2013). Motivation is inversely proportional to α, meaning a larger α value corresponds to lower motivation.

Following the baseline threshold procedure, all rats were assigned to one of three groups: Short Access (ShA; n=10), prolonged Intermittent Access (IntA; n=20), or prolonged Long Access (LgA; n=12). Groups were determined following the baseline threshold test so that there were no baseline differences between groups in α, P_Max_, or Q_0_. Following the baseline threshold procedure rats in the ShA group underwent surgery to implant a microdialysis guide cannula (described below). After three days of recovery each rat in the ShA group was habituated to the microdialysis chamber in which they would be tested for ~1 hour. Then these rats were given three more IC40 self-administration sessions and at least one of these sessions was conducted in the microdialysis chamber in which they would be tested. Thus, each rat had at least one habituation session and one self-administration session in the microdialysis chamber. Due to complications on the test day one ShA rat had to be excluded.

### Self-Administration: intermittent access or long access procedures

After their behavior stabilized on the threshold procedure, the rats in the Prolonged Access groups were given 30 self-administration sessions (Fig. 1a). These rats were trained using either an intermittent access (IntA) procedure similar to that described previously (Zimmer *et al*., 2012; Kawa *et al*., 2016) or a long access (LgA) procedure similar to that described previously (Ahmed & Koob, 1998). During the IntA procedure, the rats were placed into the chamber with the house light illuminated. The beginning of the first 5-min Drug-Available period started two minutes after the rats were placed into the chamber and was signaled by extinguishing the house light. During the Drug-Available period a nose poke into the active port resulted in an intravenous infusion of cocaine hydrochloride (NIDA) as during acquisition. Each infusion was paired with the illumination of a cue light in the nose port for the duration of the infusion. However, unlike during acquisition, rats could take another infusion immediately after the preceding one (i.e., the only time out period was during the 2.6 second infusion). After the 5-min Drug-Available period, the house light turned on and signaled a 25-min No-Drug Available period. After 25-min, the house light was extinguished and another 5-min Drug-Available period began. Each IntA session lasted 4 hours (8 cycles of Drug-Availability).

During the LgA procedure, the rats were placed into the chamber with the house light illuminated. After two minutes the house light was extinguished and the rats were able to self-administer cocaine. Each LgA session lasted 6 hours and throughout the session a nose poke into the active port resulted in an intravenous infusion of cocaine hydrochloride (NIDA) on a FR-1 schedule, as during acquisition. Each infusion was paired with the illumination of a cue light in the nose port for 20 seconds. Nose pokes during this time were recorded but had no consequences. With the exception of this timeout period, cocaine was available to the rats throughout the 6-hour session. An inactive port was also present at all times and pokes there had no consequences.

Rats in both groups underwent one self-administration session/day for an average of 5 days/week. We varied the number and pattern of days off each week to model the intermittency *between* bouts of drug use seen in human cocaine users (Cohen & Sas, 1994; Simon *et al*., 2002) - for example, one week the rats may have had only 1 day off and then the next week the rats may have had 3 days off. However, rats were never given the day directly before a probe test off. The rats were given 25 self-administration sessions and then were tested again using the within-session threshold procedure. This probe test was identical to the baseline threshold test except that the rats were only tested for two days, and the respective values were averaged across the two test days. Two rats in the IntA group were removed from testing prior to this threshold test due to loss of catheter patency. After two threshold test sessions, the rats underwent surgery to implant a microdialysis guide cannula above the nucleus accumbens core, as described below. After three days of recovery the rats were given five additional IntA or LgA self-administration sessions. Also, each rat was first habituated to the microdialysis chamber in which they would be tested for ~1 hour and then at least one of the five additional self-administration sessions was conducted in the microdialysis chamber. Thus each rat had at least one habituation session and one self-administration session in the microdialysis chamber. Then 1-3 days after the last self-administration session these rats underwent a microdialysis test and collection session identical to the ShA rats and described below. Due to complications during surgery or the microdialysis test session 2 IntA rats and 3 LgA rats were excluded, leaving 16 and 9 successful collections, respectively.

### Extinction and cue-induced reinstatement test

Following the microdialysis test session, a subset of the Prolonged Access rats was tested for the ability of the cocaine-paired discrete cue (light in the nose port) to reinstate drug-seeking (n=10 IntA; 9 LgA). The rats underwent two-hour extinction sessions (1/day) for at least 5 days and until they made less than 20 active nose pokes for two consecutive sessions. The extinction sessions consisted of placing the rats into a chamber with the house light on and the session started two minutes later. Upon the session starting, the house light turned off and remained off for the duration of the session. Responses into the nose ports during these sessions were recorded but had no consequences. The day after a rat met the extinction criterion it underwent a day of testing identical to extinction except on this day pokes in the active port were reinforced by the illumination of the cue light for 2.6-sec. After the cue-induced reinstatement test these rats were perfused to verify cannula placement.

### Microdialysis

#### Intracranial placement of cannula for microdialysis

After their behavior stabilized on the threshold procedure (ShA group) or after 25 additional self-administration sessions (IntA and LgA groups), the rats underwent surgery to implant a microdialysis guide cannula above the nucleus accumbens core. Rats were anesthetized with a ketamine xylazine cocktail as described for the catheter implantation and placed in a stereotaxic instrument (David Kopf Instruments, Tujunga, CA). A guide cannula (CMA, CMA12 Guide Cannula) was lowered such that its tip was just above the nucleus accumbens core. The coordinates were +1.6mm anterior, +/-1.6mm lateral, and -6.2mm ventral, relative to bregma (Paxinos & Watson, 2007). Hemisphere (right/left) was counter-balanced across all groups. To prevent clogging, a stainless steel stylet was inserted into the cannula. The guide cannula was secured to the skull using three metallic screws and acrylic dental cement. The rats were administered the analgesic carprofen (5 mg/kg s.c.) during their recovery from surgery. Rats were allowed at least three days of recovery before any subsequent testing.

#### Probe construction and test session

The microdialysis probes were custom made similar to those described previously (Pitchers *et al*., 2017). Briefly, two silica capillaries (75μm inner diameter; 150 μm outer diameter; TSP075150; Polymicro Technologies) were glued together and inserted into a 24-gauge stainless steel tube that served as the shaft to be inserted into the guide cannula. The portion of each capillary tube that was not inserted into the shaft was sheathed in 22-gauge stainless steel tubing and one was used for the inlet capillary and one for the outlet capillary. The capillary tip extending from the shaft was sheathed in an 18 kDa molecular weight cutoff regenerated cellulose membrane (Spectrum Labs). The tip and base of the membrane was sealed with an epoxy resin. The membrane extended 2 mm in length beyond the shaft, such that a 2 mm probe would extend into the center of the accumbens core.

In all groups, the microdialysis test session was conducted 1-3 days after the last self-administration session. Approximately 12-16 hours before the microdialysis test session, the stylet was removed from the guide cannula and a probe was inserted. The rat was placed into the microdialysis test chamber with the house light on. The probe was perfused at a rate of 0.5 μL/min with artificial cerebrospinal fluid (aCSF) overnight and the rate was increased to 1 μL/min approximately four hours prior to collection. The aCSF was comprised of 145 mM NaCl, 2.7 mM KCl, 1.0 mM MgSO_4_, 1.2 mM CaCl_2_, 1.55 mM NaHPO_4_, and 0.45 mM NaH_2_PO_4_. In addition, 100 nM ^13^C_6_-DA was added to the aCSF which allowed for *in vivo* calibration of the probes (Hershey & Kennedy, 2013) in all but a small subset of the rats (3 ShA, 4 IntA). In this subset of rats recovery rates for each probe were calculated by placing the membrane in known concentrations of DA and correcting for the concentration collected in the dialysate sample. These rats did not differ on any measures from rats tested with ^13^C_6_-DA added to the aCSF and thus were combined for analysis. Approximately four hours prior to collection the rats were also attached to the drug delivery tubing. Dialysate samples were collected every 3 minutes.

The test session and collection started with ten baseline samples (30 minutes). After 30 minutes, the house light turned off which signaled that cocaine was available, as it had during all of the previous self-administration sessions. No samples were collected until the rat had made one response in the active nose port and self-administered a single 1.25 mg/kg cocaine infusion. The light cue that had been paired with cocaine infusions during self-administration testing was *not* presented, to better isolate the effect of cocaine alone. As soon as the rat made a response and the infusion ended the house light turned back on and no additional cocaine was available to the rat. After accounting for the time it took for the dialysate to travel from the brain to the outlet collection, dialysis samples were collected for 1 hour (20 samples) that corresponded to the onset of the cocaine infusion. One hour after the cocaine infusion, the house light again turned off and the light in the active nose port that had previously been a conditioned stimulus paired with cocaine delivery during self-administration was flashed 14 times for 2.6 seconds per flash over a three minute period. We collected a single sample that corresponded to this three-minute cue-presentation and five more samples following the cue presentation. Nose pokes in both the active and inactive nose ports were recorded throughout the test session. In summary, 3-min samples were collected over a 30-minute baseline period, for 1-hour following a cocaine infusion, for three minutes during which the cue was presented non-contingently, and then for 15 minutes following cue presentation.

#### Analysis of neurochemical levels in dialysate samples using high performance liquid chromatography coupled with mass spectrometry (HPLC-MS)

All reagents, drugs, and chemicals were purchased from Sigma-Aldrich (St. Louis, MO) unless otherwise noted. Samples were analyzed using benzoyl chloride derivatization and a modified LC-MS method previously described (Song *et al*., 2012). Briefly, the 3 μL samples were derivatized by adding 1.5 μL of 100 mM sodium carbonate monohydrate buffer, 1.5 μL of 2% benzoyl chloride in acetonitrile, and 1.5 μL of an internal standard mixture (to improve quantification), in order, briefly vortexing between each addition. A Thermo Fisher (Waltham, MA) Vanquish UHPLC system automatically injected 5 μL of the sample onto a Phenomenex Kinetex C18 HPLC column (2.1mm X 100mm, 1.7 μm). Mobile phase A consisted of 10 mM ammonium formate and 0.15% formic acid. Mobile phase B was pure acetonitrile. Analytes were detected using positive electrospray ionization with a Thermo Fisher TSQ Quantum Ultra triple-quadrupole mass spectrometer operating in multiple-reaction monitoring (MRM) mode.

Inclusion of ^13^C_6_ dopamine in the aCSF perfusate allowed us to calculate an extraction fraction (E_d_) for each sample (Hershey & Kennedy, 2013). The E_d_ (E_d_=1-(C_in_/C_out_) is the ratio of the amount of the isotope that exits the probe to the amount retained in the dialysate sample. Absolute extracellular concentrations of dopamine were determined by dividing the dialysate concentration of dopamine by the E_d_ value. Finally, in a subset of rats we switched the perfusate to aCSF that lacked ^13^C_6_ dopamine at the conclusion of the test session. After allowing this aCSF to run for ~one hour we collected several samples in the same manner as the samples collected during the test. We analyzed these samples and compared them to samples from the baseline portion of the test session. This analysis, along with comparisons between rats that did not have ^13^C_6_-DA added to the aCSF at any point, confirmed that there were no effects of infusing 100 nM ^13^C_6_-DA on endogenous dopamine levels (Fig. 3a).

### Histological Analysis

After the microdialysis test session, rats were anesthetized using sodium pentobarbital (270 mg/kg; i.p.) and perfused intracardially with 50 mL of 0.9% saline, followed by 500 mL of 4% paraformaldehyde in 0.1 M phosphate buffer (PB). After being perfused, brains were removed, post-fixed in the same paraformaldehyde solution for 2 hours, then immersed in 20% sucrose and 0.01% sodium azide in 0.1 M PB for 48 hours at 4° C. Coronal sections (40 μm) were cut with a freezing microtome (SM 2000R; Leica), collected in PB, and mounted on to a slide immediately. Sections were imaged at 4x magnification using a Leica DM400B digital microscope to verify cannula placement.

### Statistical analysis

Linear mixed-models (LMM) analyses were used for all behavioral repeated measures data. The best-fitting model of repeated measures covariance was determined by the lowest Akaike information criterion score (West *et al*., 2007). Depending on the model selected, the degrees of freedom may have been adjusted to a non-integer value. Data for the α measure was not normally distributed and therefore all statistical tests involving α were run on log transformed data, consistent with previous reports (Bentzley *et al*., 2014). Planned post-hoc contrasts (and Bonferroni corrections) were done to compare between the different self-administration procedures and within the Prolonged groups across the two test periods. Active and inactive nose pokes from the cue-induced reinstatement test were compared to the last day of extinction and between the IntA and LgA groups. Similar LMM analysis and planned post-hoc analysis were used to analyze neurochemical levels from the microdialysis test session. Statistical significance was set at p<0.05.

## RESULTS

### LgA produced more cocaine consumption than IntA, but rats in both groups escalated their intake to a similar extent

The overall timeline of the experiment is shown in Fig. 1a. Fig. 1b shows the acquisition of cocaine self-administration as a function of Infusion Criterion for all rats, prior to group assignment. The number of responses at the active nose port increased across sessions (effect of IC, F(2,40.4)=38.7, p<0.001), and the number of responses at the inactive nose port decreased across sessions (effect of IC, F(2,75.1)=3.5, p=0.04). There was no difference between the groups on any measure of acquisition.

A subset of rats was given 30 self-administration sessions using either the IntA or LgA procedure. Rats in the IntA group quickly learned to discriminate between the Drug-Available and No-Drug periods, as previously reported (data not shown; see Kawa *et al*., 2016). When cocaine intake per session from the IntA and LgA rats was analyzed together there was a main effect of test session (F(29,70.7)=4.8, p<0.001; Fig. 1c). Further analysis confirmed that when each group was analyzed separately both LgA (p=0.02; Fig. 1c inset) and IntA (p=0.01, Fig. 1c inset) rats escalated their intake as a function of increasing self-administration experience. Relatedly, the rate at which consumption increased did not differ between the groups (group X session interaction, F(29,70.7)=1.2, p=0.24). There was however a large group difference in the total amount of cocaine consumed (effect of group, F(1,92.3)=823, p<0.001; Fig. 1d). LgA rats consumed ~2.8x more cocaine than IntA rats.

### Prolonged IntA and LgA experience had different effects on motivated behavior

All rats underwent a baseline threshold test to quantify their motivation to self-administer cocaine following only limited self-administration experience (Fig. 2). Group assignments (ShA, IntA, LgA) were made following this test, such that there were no differences between the groups on any of the measures derived from this test. Accordingly, the baseline values shown in Figure 2 are averages that include all rats, for illustrative purposes. The rats that were placed into the IntA and LgA groups then underwent 25 self-administration sessions before another threshold probe test to determine how motivation changed as a function of IntA or LgA experience, relative to baseline. All statistical comparisons to baseline were within-subject.

**Fig. 2.**
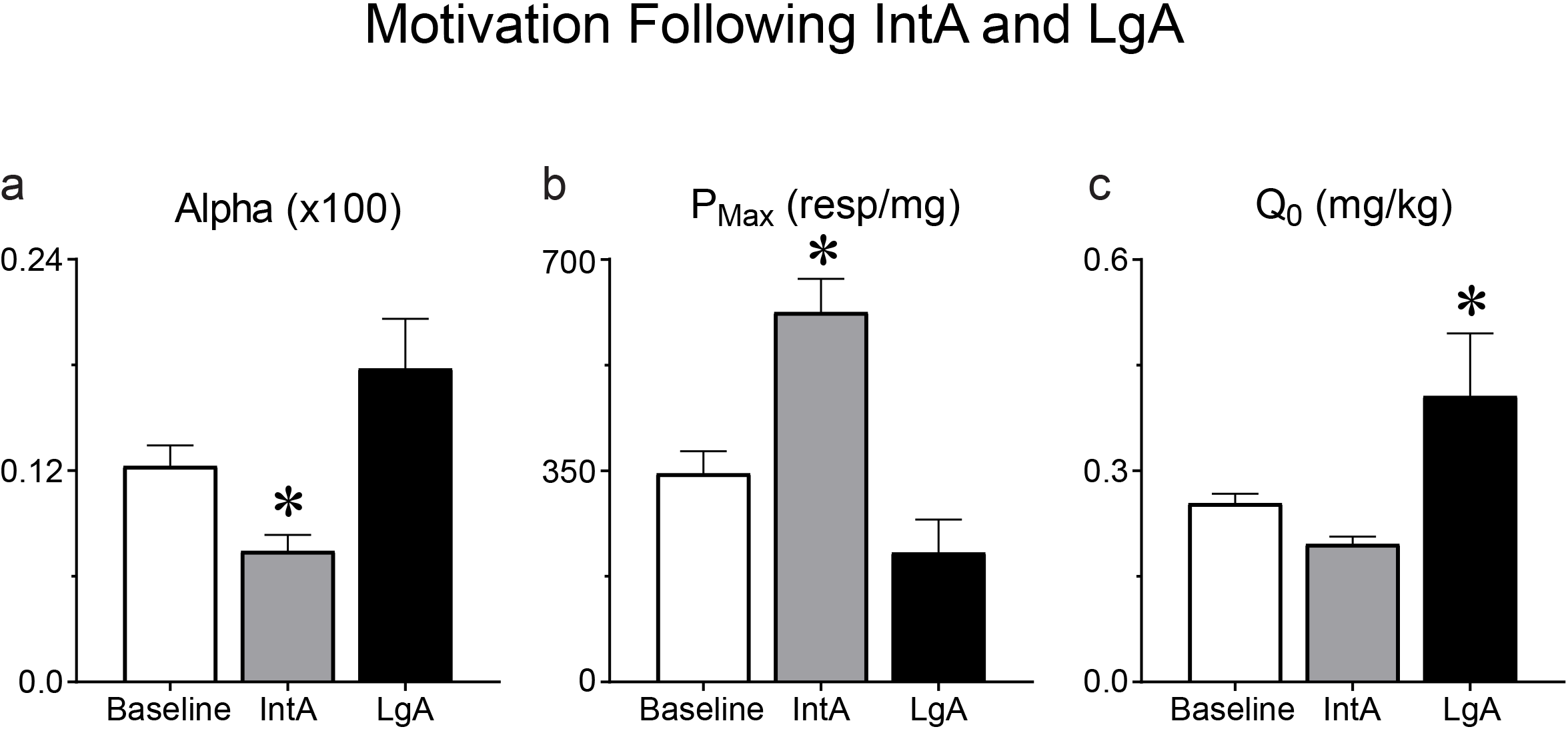
Cocaine demand was assessed using a threshold procedure in rats with different self-administration experience. In panels a-c all rats are included in the ‘Baseline’ group (n=42) and a subset of rats went on to experience IntA self-administration (n=18) while a separate group of rats went on to LgA self-administration (n=12). IntA experience decreased alpha (α; increased motivation) relative to LgA experience (**a**) and increased P_Max_ (increased motivation) relative to LgA experience (**b**). LgA experience increased Q_0_ (preferred cocaine intake when effort is not a factor) relative to IntA experience (**c**). Values represent means ± SEMs, * represents a significant difference (p<0.05) between IntA rats and LgA rats.

The metric α measures the elasticity of the demand curve generated during the within-session threshold procedure, and is inversely proportional to motivation (Bentzley *et al*., 2013; Fig. 2a). Prolonged IntA and LgA experience had different effects on α, as indicated by a significant group X probe test interaction (F(1,69)=7.2, p=0.009). Post-hoc within-subject analysis revealed that IntA experience decreased α (increased motivation) (p=0.02) whereas LgA experience did not significantly change α (p=0.12). On the second probe test IntA rats had a lower α than LgA rats (effect of group, p=0.04), indicating that rats with IntA experience were more motivated to obtain cocaine.

The metric P_Max_ is a measure of motivation that reflects the maximum price an individual is willing to pay (in effort) to obtain a reinforcer (Bentzley *et al*., 2013; Fig. 2b). Again, prolonged IntA and LgA had different effects on P_Max_, as indicated by a significant group X probe test interaction (F(1,33.1)=10.4, p=0.003). Post-hoc within-subject analysis revealed that IntA experience increased P_Max_ (p=0.001), and LgA experience did not significantly change P_Max_ (p=0.27). On the second probe test IntA rats had a higher P_Max_ than LgA rats (effect of group, p=0.005), indicating that they were more motivated to obtain cocaine, consistent with α.

Another metric derived from the threshold procedure is Q_0_, which measures the preferred level of cocaine consumption when cost is nil (Bentzley *et al*., 2013; Fig. 2c). Again there was a significant group X probe test interaction (F(1,28.7)=7.7, p=0.009) indicating that IntA and LgA had different effects on Q_0_. Q_0_ differed from α and P_Max_ as post-hoc within-subject analysis revealed that LgA experience increased Q_0_ (p=0.01) but IntA experience had no effect on Q_0_ (p=0.25). Finally, LgA rats had higher Q_0_ than IntA rats after prolonged self-administration (effect of group, p=0.03). These differential effects of LgA and IntA on Q_0_ are consistent with several reports (Oleson & Roberts, 2009; Bentzley *et al*., 2014; Kawa *et al*., 2016; James *et al*., 2018; Singer *et al*., 2018).

### Infusion of stable-isotope labeled dopamine did not affect endogenous dopamine levels

Use of extraction fraction (E_d_) to calculate *in vivo* concentration accounts for possible differences in recovery associated with the probe and brain environment in different rats. *In vivo* calibration therefore improves the accuracy of such measures and allows comparison between rats. 100 nM stable-isotope labeled dopamine (^13^C_6_-DA) was included in the aCSF that was perfused during the microdialysis test session to calculate an E_d_ for each sample (see methods; Hershey & Kennedy, 2013). As a control, we determined if the infusion of this concentration of ^13^C_6_-DA affected endogenous DA levels. For this control measurement, the input solution was switched to aCSF that lacked ^13^C_6_-DA at the end of each dialysis test session and after allowing the lines to clear of any ^13^C6-DA aCSF three more dialysate samples were collected to compare to the last samples collected with ^13^C_6_-DA aCSF. As we have reported previously (Hershey & Kennedy, 2013) the presence of ^13^C_6_-DA did not affect endogenous DA levels (Fig. 3a). In addition, the calculated E_d_ did not differ between groups nor did they change across the dialysis test session (all p-values>0.1; Fig. 3b). These results provide confidence that the calibrated DA concentrations reported can be compared between rats and groups.

**Fig. 3.**
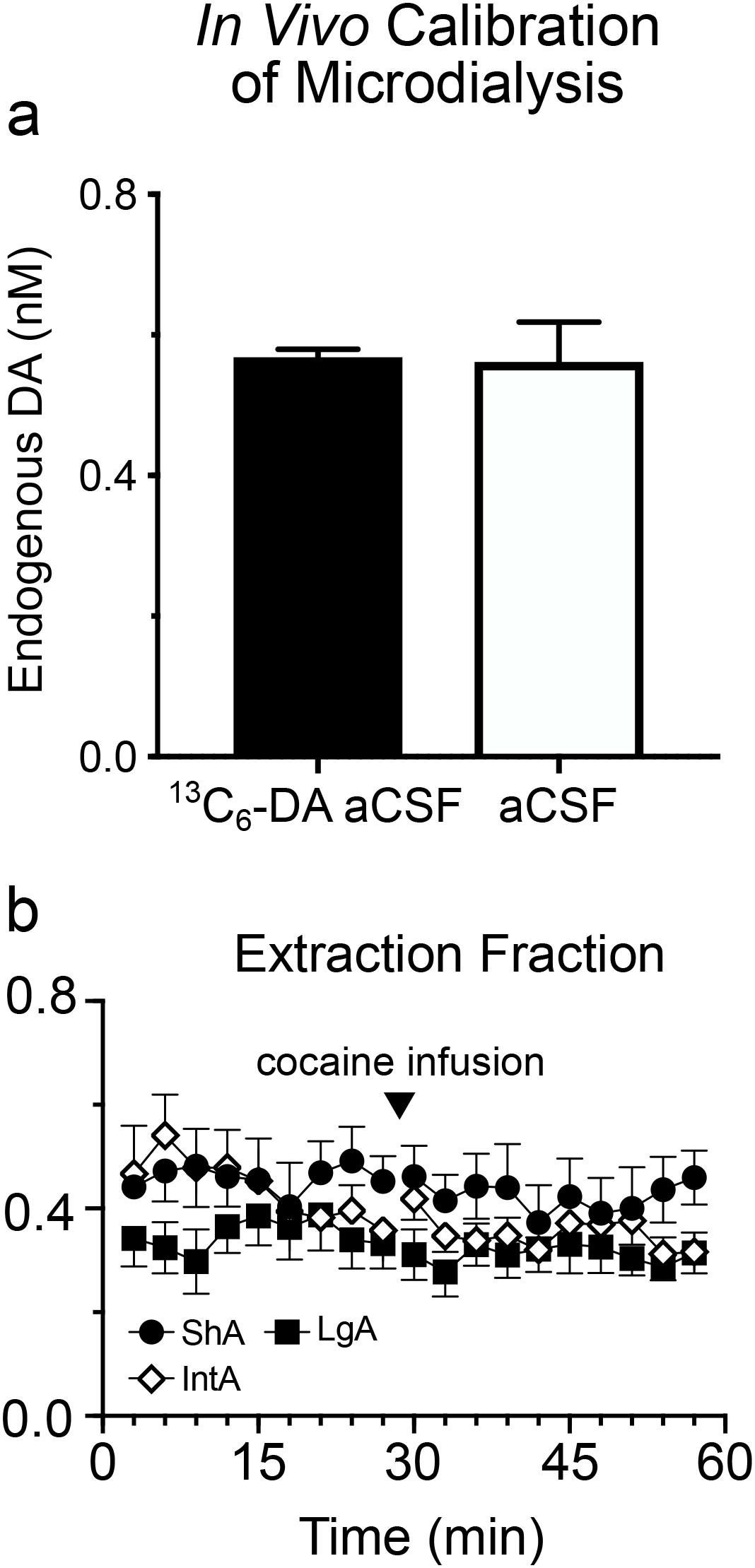
Stable-isotope labeled DA (^13^C_6_-DA) was included in the aCSF perfusate during the dialysis test session in order to calculate an extraction fraction (E_d_) for each sample collected. Infusing ^13^C_6_-DA did not affect endogenous DA levels (here not corrected for probe recovery) when compared to aCSF that lacked ^13^C_6_-DA (**a**). The average calculated E_d_ throughout the test did not differ between ShA, LgA, or IntA rats (10 baseline and 10 post-cocaine samples shown) (**b**). Values represent means ± SEMs.

### Prolonged IntA (but not LgA) sensitized cocaine-induced dopamine overflow

Following the collection of baseline dialysate for 30 min, rats with limited ShA, prolonged IntA, or prolonged LgA experience were allowed to self-administer a single IV injection of 1.25 mg/kg cocaine (see Fig. 4a for timeline), and dialysate was then collected for an additional 60 min. There were no group differences in the latency to self-administer the cocaine infusion as most rats nose poked as soon as the house light turned off signaling drug availability. We first analyzed the average concentration of DA in the ten baseline samples and the first ten post-cocaine samples in all 3 groups (Fig. 4b). When analyzing all 20 samples, there was a main effect of group (F(2,326)=10.2, p<0.001) and the cocaine infusion increased extracellular DA, relative to baseline, in all groups (main effect of cocaine, F(1,331)=8.93, p=0.003; Fig. 4b). Planned post-hoc analysis revealed that the main effect of group was due to a differential response to the cocaine infusion. That is, there were no group differences in baseline DA levels (all p-values>0.1) but following the cocaine infusion, DA levels increased to a greater extent in IntA rats, relative to LgA rats (p<0.001) and ShA rats (p=0.004). The latter two groups did not differ (p=0.21). The largest DA response to cocaine reliably occurred within the first three samples (9 minutes) after the infusion, but there was individual variation in the time that elapsed from infusion to peak response. To better visualize the peak response to cocaine in all individuals Figure 4c shows the DA response in all three groups when each rat’s peak DA response was aligned.

**Fig. 4.**
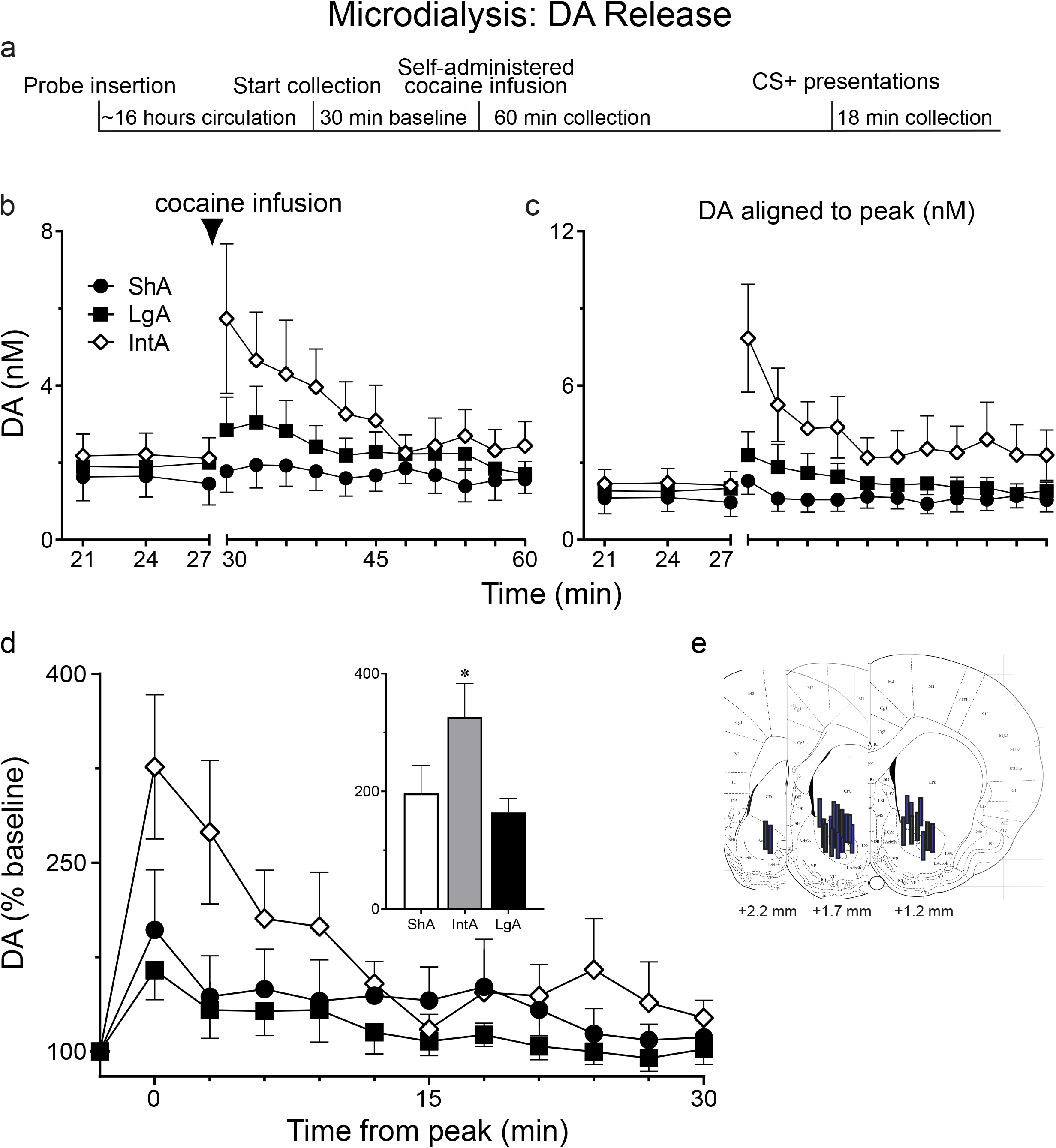
Dopamine (DA) overflow in response to a self-administered cocaine infusion was tested in rats with different self-administration experience. A timeline of the microdialysis test day is shown in panel **a**. Data from the conditioned stimulus (CS presentation) is not shown as there was no change in DA levels. Cocaine-evoked DA overflow was greater in rats trained with IntA (n=16) than rats trained with LgA (n=9) or ShA (n=9) (**b**). Panel **c** shows the same data with each rat’s peak response to cocaine aligned. When peak DA release was analyzed as a percent of baseline, cocaine evoked a larger DA response in IntA rats than LgA or ShA rats (**d**). Microdialysis probes were targeted to the nucleus accumbens core (**e**). Values represent means ± SEMs.

Given that there were no group differences in baseline DA levels, we averaged the baseline values together for each group to determine the percent change from baseline produced by cocaine (Fig. 4d). For this analysis we aligned each rat’s peak DA response. There was a main effect of group (F(2,672)=6.04, p=0.003) and the cocaine infusion increased extracellular DA in all groups (F(1,673)=40, p<0.001). Further, the cocaine infusion increased DA levels to a different extent in the three groups evidenced by a group X cocaine interaction (F(2,672)=6.04, p=0.003). Planned post-hoc analysis revealed that the cocaine infusion increased DA levels to a greater extent in IntA rats than LgA rats (p<0.001) or ShA rats (p=0.016), and the latter two groups did not differ (p=0.1), consistent with the analysis of extracellular DA that was not normalized to baseline.

Active nose pokes that were made in the hour following the single self-administered cocaine infusion during the microdialysis test session were recorded and analyzed as a measure of cocaine-induced drug seeking. A one-way Anova comparing the three groups revealed no effect of group on nose pokes, although there was a trend towards IntA rats making more responses than ShA or LgA rats (effect of group, F(2)=2.7, p=0.08; data not shown).

### IntA rats showed greater cue-induced reinstatement than LgA rats

The day after the microdialysis test session, a subset of the rats in the IntA group and the LgA groups underwent extinction training followed by a test for cue-induced reinstatement of cocaine-seeking (conditioned reinforcement). There were no group differences during extinction training in the number of responses made or the number of sessions required to reach extinction criteria (all p-values>0.1; Fig. 5a). On the test day, when responding was reinforced by the presentation of the cue that had previously been paired with cocaine, both IntA and LgA rats reinstated their drug-seeking relative to extinction levels (F(1,20.1)=100, p<0.001; Fig. 5b), specifically at the active nose port (effect of inactive nose port X session, F(1,18.1)=2.3, p=0.15). However, rats with IntA experience showed more robust cue-induced reinstatement than rats with LgA experience (effect of group, F(1,20.1)=22, p<0.001).

**Fig. 5.**
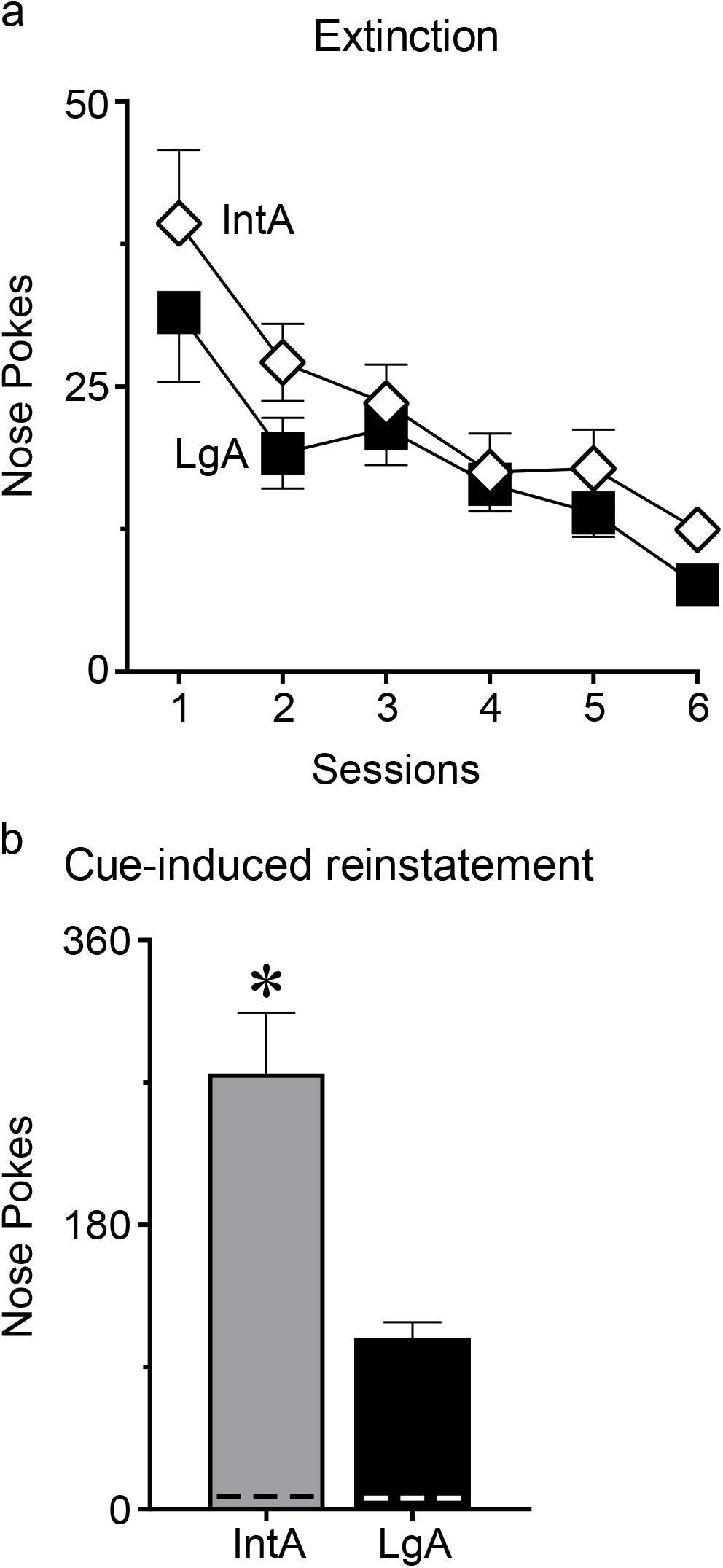
After the dialysis test session a subset of IntA rats (n=10) and LgA rats (n=9) were tested for the ability of a cocaine-paired cue to reinstate drug-seeking. IntA and LgA rats did not differ in responding during extinction training (**a**). In a test for conditioned reinforcement, when a nose poke in the previously active port was reinforced by presentation of the cue that had previously been associated with cocaine but not cocaine itself, IntA rats made more responses than LgA rats (**b**). Nose pokes at the inactive port are represented by dashed lines. Values represent means ± SEMs, * represents a significant difference (p<0.05) between IntA rats and LgA rats.

### Cocaine-seeking on multiple tests of addiction-like behavior predicted cocaine-evoked dopamine overflow

Given the role of DA in motivated behavior we asked whether DA overflow in response to self-administered cocaine correlated with cocaine-seeking behavior and/or motivation for cocaine. Fig. 6 shows that the magnitude of the increase in DA, as a percent of baseline, on the microdialysis test day was positively correlated with cocaine-seeking on the dialysis test day, assessed by the number of nose pokes into the active port (R^2^=0.14, p=0.03; Fig. 6a). There was a similar correlation between cocaine-induced DA and motivation for cocaine (P_Max_) assessed on the most recent threshold test (first test for ShA; second for IntA/LgA) (R^2^=0.32, p<0.001; Fig. 6b). This correlation was evident even when the analysis was restricted to the IntA group (R^2^ =0 .29, p=0.03). There was a similar correlation between the DA response and α (R^2^ =0.13, p=0.04; data not shown). In contrast, Q_0_ did not correlate with the DA response when all groups were combined (R^2^=0.07, p>0.1; Fig. 6c) or within any individual group (all p-values>0.1).

**Fig. 6.**
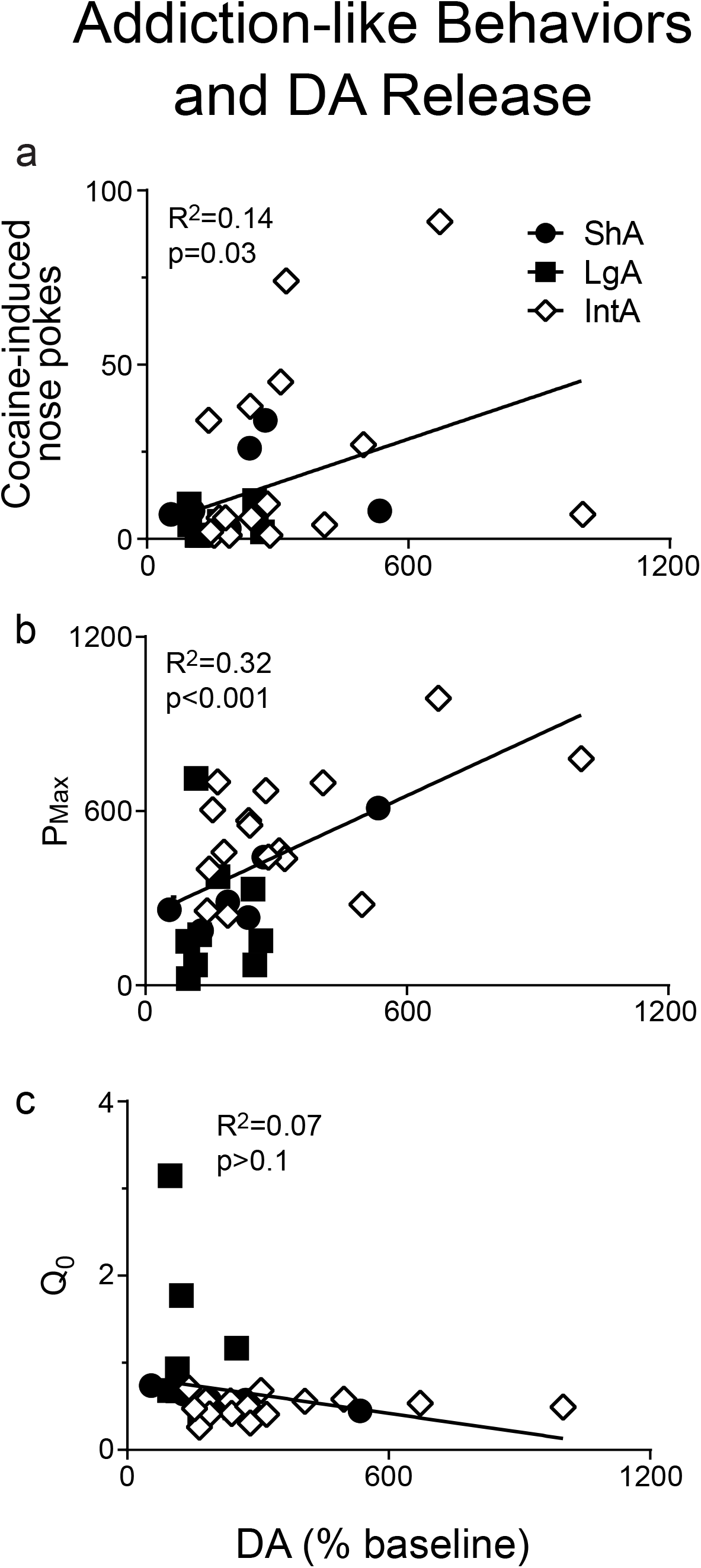
Each rat’s peak dopamine (DA) response to a self-administered cocaine infusion, as a percent of baseline, was correlated with that rat’s performance on several tests of addiction-like behavior. The peak DA change, as a percent of baseline, predicted cocaine-induced nose pokes (measured during the microdialysis test session) (**a**) and P_Max_ from the most recent threshold test (**b**). Peak DA release, as a percent of baseline, did not predict Q_0_ from the most recent threshold test (**c**).

IntA experience reliably produces multiple addiction-like behaviors and this has been shown to be particularly robust in susceptible individuals (Kawa *et al*., 2016; Singer *et al*., 2018). Thus, we separated rats that were trained with the IntA procedure into those that met 2/3 ‘addiction criteria’ and those that met 0/1 ‘addiction criteria’, as described previously (Deroche-Gamonet *et al*., 2004; Kawa *et al*., 2016; Singer *et al*., 2018). Briefly, a rat met a criterion for addiction if it was within the top third of the population on a given measure. The measures used here were α, nose pokes during the No-Drug period of IntA, and cocaine-induced nose pokes during the microdialysis test session (Fig. 7). There were 9 rats that met 0/1 criteria and 7 rats that met 2/3 criteria. Not surprisingly, 0/1 and 2/3 criteria rats differed in their performance on the behavioral measures for which they were classified (Fig. 7 a, b & c). Importantly, 2/3 criteria rats showed greater cocaine-evoked DA release than 0/1 criteria rats (p=0.05; Fig. 7d). Finally, when 2/3 criteria IntA rats (the most “addict-like”) were compared to ShA and LgA rats (that largely did not show addiction-like behavior), the cocaine infusion increased DA to a greater extent in 2/3 criteria IntA rats (group X cocaine interaction, F(2,484)=19.5, p<0.001; Fig. 7e).

**Fig. 7.**
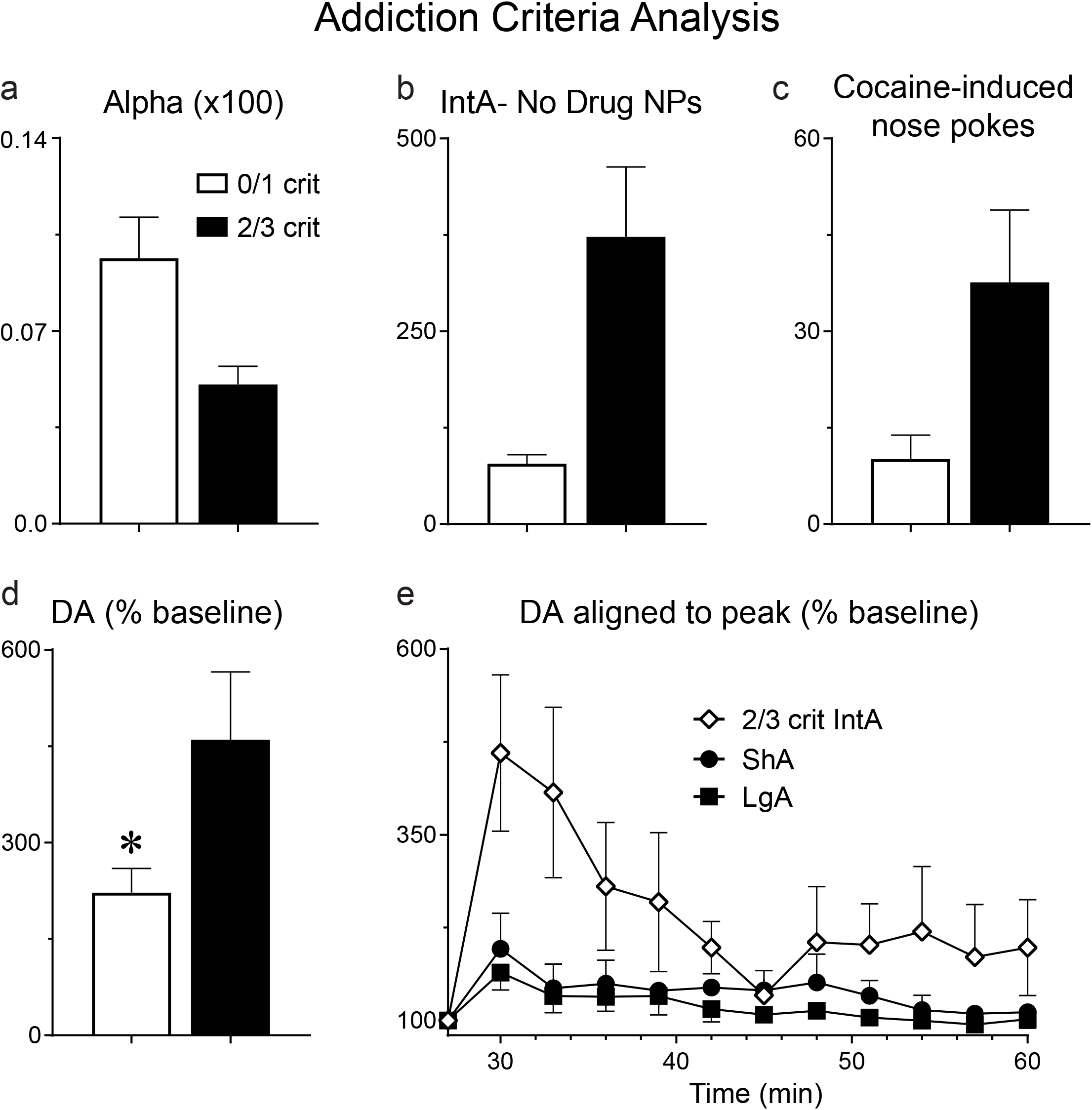
Analysis based on addiction criteria. IntA rats were separated based on the number of ‘addiction-criteria’ they met (see methods). Rats that met 2/3 criteria (n=7) showed increased motivation relative to rats that met 0/1 criteria (n=9) as indicated by alpha (α) (**a**), nose pokes (**NPs**) made during the No Drug periods of IntA (**b**), and cocaine-induced nose pokes (measured during the microdialysis test day) (**c**). A self-administered infusion of cocaine evoked greater peak dopamine (DA) release, shown as a percent of baseline, in 2/3 criteria rats than 0/1 criteria rats (**d**). Panel **e** shows the timecourse of DA release (% baseline) following cocaine in 2/3 criteria IntA rats, ShA rats, and LgA rats. Values represent means ± SEMs.

### Self-administered cocaine increased extracellular glutamate and 3-MT levels

The HPLC-MS technique used here allows for the analysis of a number of analytes, in addition to DA (Table 1). There were no group differences in the baseline levels of any of the analytes quantified. Therefore, all values were normalized to baseline and cocaine-induced changes relative to baseline were calculated. Self-administered cocaine increased extracellular glutamate levels relative to baseline in all groups (effect of cocaine, F(1,671)=3.97, p=0.04), but there were no group differences (p>0.1). In addition, self-administered cocaine increased 3-MT, a DA metabolite formed extracellularly, in all groups (effect of cocaine, F(1,581)=10.5, p<0.001). Further, cocaine increased 3-MT to a greater extent in IntA rats than LgA or ShA rats (group X cocaine interaction, F(2,580)=4.93, p=0.008; post-hoc analysis revealed this to be driven by IntA rats). We also analyzed extracellular GABA, ACh, DOPAC, and HVA levels but none of these differed between groups or changed significantly following cocaine.

**Table 1.**
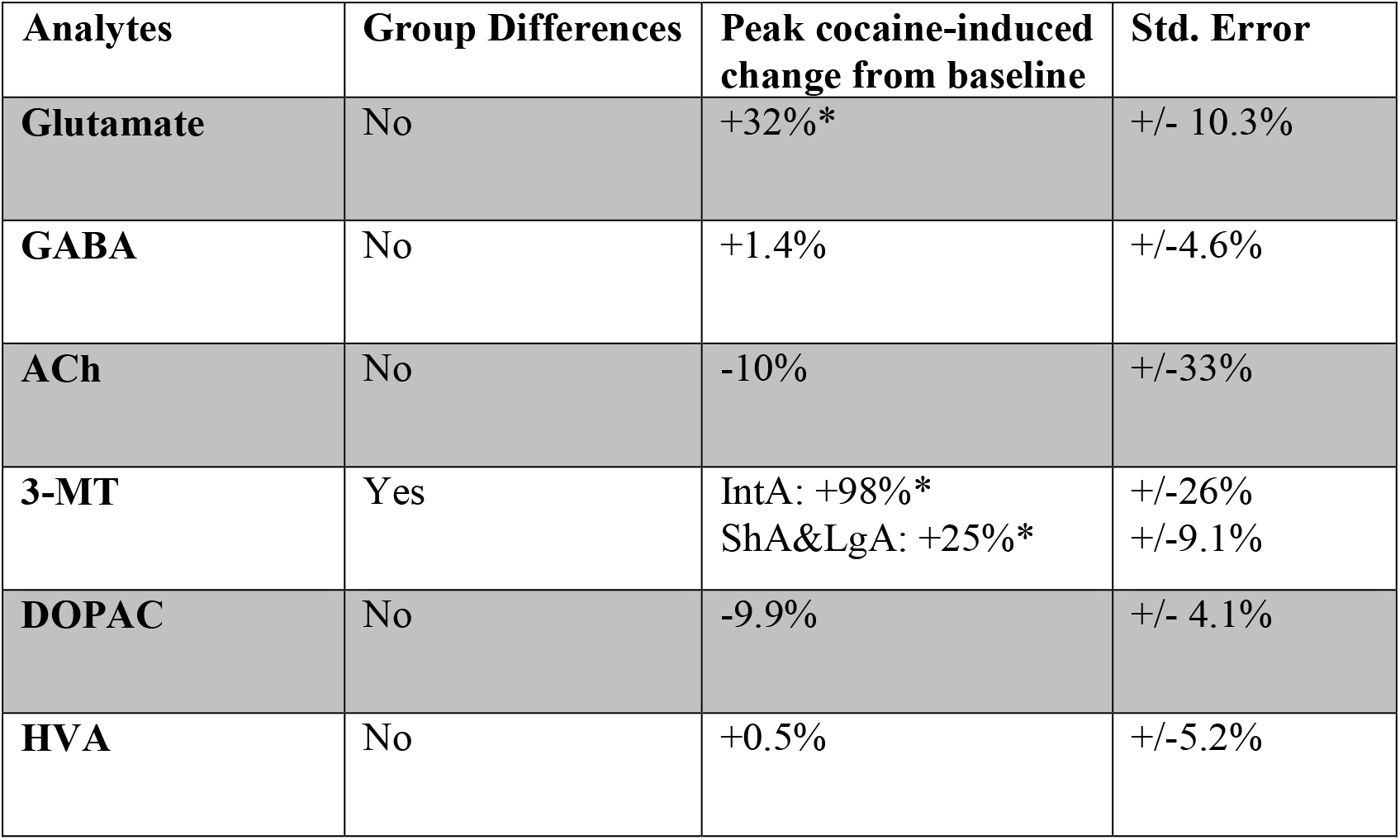
Analytes of interest following a cocaine infusion. Microdialysis allows for the analysis of a number of analytes in addition to DA. Select neurochemicals of interest are shown here. A self-administered cocaine infusion increased extracellular levels of Glutamate and 3-MT (a dopamine metabolite) in all rats. Cocaine-evoked 3-MT was greater in IntA rats than ShA rats or LgA rats. * indicates a significant (p<0.05) change in analyte concentration following the cocaine infusion relative to baseline.

### Presentation of the cocaine-paired cue did not affect extracellular neurochemical levels

One hour after the self-administered cocaine infusion, the conditioned stimulus that had been paired with cocaine infusions during self-administration training was presented non-contingently for three minutes. Presentation of the conditioned stimulus did not produce any detectable changes in any of the analytes that we measured (data not shown; see discussion).

## DISCUSSION

The purpose of this study was to compare the ability of prolonged LgA and IntA cocaine self-administration experience to produce addiction-like behavior (relative to ShA), and how this influenced the ability of self-administered cocaine to change extracellular DA levels in the nucleus accumbens core *in vivo*. The main findings were: 1. As expected, LgA resulted in much greater total cocaine consumption than IntA. 2. Both IntA and LgA produced escalation of intake with increasing self-administration experience. 3. IntA (but not LgA) experience increased motivation for cocaine, as indicated by a decrease in α and increase in P_Max_. 4. IntA rats showed greater cue-induced reinstatement of cocaine-seeking than LgA rats. 5. LgA (but not IntA) experience increased the preferred level of cocaine intake when no effort was required (Q_0_). 6. There were no group differences in the basal levels of DA in dialysate, but a single self-administered IV injection of cocaine increased DA in the core of the nucleus accumbens to a greater extent in rats with prior IntA experience than those with LgA or ShA experience, and the latter two groups did not differ. 7. Across all groups high motivation for cocaine was associated with a greater DA response. 8. There were no group differences in dialysate concentrations of glutamate, GABA, ACh, DOPAC or HVA, although cocaine increased 3-MT to a greater extent in IntA than ShA or LgA rats, consistent with the effects on DA.

### IntA experience was more effective in producing addiction-like behavior than LgA experience

Since its introduction in 1998 (Ahmed & Koob, 1998) the LgA procedure has been widely adopted to model the transition to cocaine addiction in rats, because it was thought to be especially effective in producing a number of addiction-like behaviors, relative to ShA (for reviews see, Ahmed, 2012; Edwards and Koob 2013). In their 1998 paper Ahmed and Koob reported that LgA, but not ShA, resulted in escalation of intake. Since that time it has also been reported that, relative to ShA, rats with LgA experience are more motivated to seek cocaine (Paterson & Markou, 2003; Wee *et al*., 2008), take more cocaine in the face of adverse consequences (Xue *et al*., 2012; Bentzley *et al*., 2014; also see Vanderschuren & Everitt, 2004), and show greater reinstatement of cocaine-seeking behavior following extinction (Mantsch *et al*., 2004, 2008; Ahmed & Cador, 2006; Kippin *et al*., 2006). As indicated by the excerpt cited in the Introduction from Ahmed (2012), it has been suggested that the critical factor necessary for the emergence of escalation and other addiction-like behavior is the *amount* of drug consumed. As put by Edwards and Koob (2013), “excessive drug exposure likely remains an indispensable element driving the development of addiction”. However, the findings presented here add to a growing literature indicating that this is not the case.

IntA self-administration results in much less total cocaine consumption than LgA. Yet as reported here, IntA also produced escalation of intake and was more effective than LgA in increasing motivation for cocaine and in producing cue-induced reinstatement of cocaine-seeking behavior. These findings are consistent with a number of recent studies that also report IntA produces escalation of intake, heightened motivation for cocaine, continued cocaine-seeking in the face of an adverse consequence, continued cocaine-seeking when it is not available, and greater cue-induced reinstatement (Zimmer *et al*., 2012; Kawa *et al*., 2016; Allain & Samaha, 2018; Allain *et al*., 2018; James *et al*., 2018; Kawa & Robinson, 2018; Singer *et al*., 2018). Collectively, these studies have established that the consumption of the large amount of cocaine associated with LgA is not necessary for the development of addiction-like behavior and other pharmacokinetic factors appear to be more important (Allain *et al*., 2015).

The failure of LgA experience to increase motivation for cocaine in the present study is inconsistent with several previous studies using either the same behavioral economic indicators used here (Zimmer *et al*., 2012; Bentzley *et al*., 2014) or Progressive Ratio (PR) tests (Paterson & Markou, 2003; Wee *et al*., 2008). However, the effects of LgA reported in these studies were often only assessed at a single time point and compared to ShA, and did not involve within-subject comparisons. In the studies that measured how motivation *changed* with increasing LgA experience (Bentzley *et al*., 2014) the effects were modest relative to the changes that occur following IntA. Our findings are consistent with other reports that LgA experience does not increase motivation for cocaine, as assessed with either behavioral economic metrics (Oleson & Roberts, 2009) or PR tests (Liu *et al*., 2005; Quadros & Miczek, 2009; Willuhn *et al*., 2014 supplementary). In addition, it has been reported that changes in motivation produced by LgA experience are very transient, lasting for only a few days after the last self-administration session (Bentzley *et al*., 2014; James *et al*., 2018), whereas the increased motivation produced by IntA experience is long-lasting - still evident after 50 days of abstinence (James *et al*., 2018). In summary, evidence that LgA increases motivation for cocaine is somewhat mixed, whereas IntA has been consistently reported to do so.

When allowed to self-administer cocaine under low Fixed Ratio (FR) schedules of reinforcement, rats generally titrate their responding to achieve a preferred brain concentration of cocaine, which they defend within a wide range of doses (Gerber & Wise, 1989; Ahmed & Koob, 1999; Lynch & Carroll, 2001). This preferred level of consumption was quantified here by the metric Q_0_ - the preferred level of consumption when cost is nil. Q_0_ presumably represents the brain level of cocaine that produces some optimal desired effect, such that neither more nor less cocaine is better. Some have referred to Q_0_ as a “hedonic set-point” (Bentzley *et al*., 2013), although “settling-point” may be more appropriate (see Berridge, 2004). Of course, it is not possible to know if Q_0_ actually reflects subjective hedonic effects in rodents. Nevertheless, LgA experience does increase the preferred level of cocaine consumption, as indicated by escalation of intake (Ahmed & Koob, 1998), and by an increase in Q_0_, as reported here and by others (Oleson & Roberts, 2009; Bentzley *et al*., 2014; James *et al*., 2018). The present results suggest, therefore, that LgA experience produces tolerance to whatever desired effect of cocaine is defended as price increases, without any change in motivation for cocaine. In contrast, IntA increases motivation for cocaine without any concomitent change in cocaine’s desired effects. Although highly speculative, this may reflect a dissociation between cocaine “wanting” and “liking” (Robinson & Berridge, 1993; Berridge & Robinson, 2016). It also suggests that cocaine intake may escalate with both LgA and IntA, but for very different reasons - because of tolerance to cocaine’s desired effect in the case of LgA and incentive-sensitization in the case of IntA (Kawa *et al*., 2016; Kawa & Robinson, 2018). Of course, the idea that consummatory and motivational aspects of behavior are psychologically (and neurobiologically) dissociable has been suggested frequently (e.g., Nicola & Deadwyler, 2000; Sharpe & Samson, 2001; Oleson *et al*., 2011; Guillem *et al*., 2014).

### Neither LgA nor IntA experience altered basal dopamine

A decrease in basal DA levels has been reported when testing occurred soon after the discontinuation of high-dose and/or high-intake cocaine self-administration procedures (Mateo *et al*., 2005; Ferris *et al*., 2011). However, in the present study neither LgA nor IntA experience had any effect on basal DA in dialysate. Also, we included ^13^C6 dopamine in the aCSF, which allowed us to calculate an extraction fraction for each sample, and thus more accurately estimate basal DA. There were no group differences in the extraction fraction, thus bolstering our conclusion that neither LgA nor IntA changed basal DA (relative to ShA). This result is consistent with other reports that LgA experience does not alter baseline DA concentrations in dialysate, relative to ShA rats (Ahmed *et al*., 2003) or drug-naïve rats (Calipari *et al*., 2014). In addition, baseline DA levels did not correlate with any of our measures of addiction-like behavior, consistent with other studies (Hurd *et al*., 1989; Ahmed *et al*., 2003).

### IntA, but not LgA, sensitizes cocaine-evoked dopamine overflow

There have been very few studies on the neurobiological consequences of IntA experience, and those available have all involved *ex vivo* measures. Most relevant to the present study are reports by Calipari et al. (2013, 2015) that IntA experience sensitizes stimulated DA release from the nucleus accumbens core in tissue slices, relative to naïve rats or rats with a history of ShA, and also increases the ability of cocaine to inhibit DA uptake. The main purpose of the present experiment was to determine if a similar sensitization of DA neurotransmission is present in awake, behaving rats. Following prolonged IntA experience a single self-administered cocaine infusion, given in the absence of the cocaine cue, produced a greater increase in extracellular DA in the accumbens core than following either LgA or ShA experience, and these latter two groups did not differ. Furthermore, the magnitude of the DA response predicted motivation for cocaine, as assessed by a number of measures, including P_Max_, α, and cocaine-seeking on the microdialysis test day. In addition, the DA response to cocaine was greatest in rats that met the most criteria for addiction. These findings establish that IntA, a cocaine self-administration procedure that is especially effective in producing incentive-sensitization and addiction-like behavior, also sensitizes the dopaminergic response to cocaine. Finally, IntA also has been reported to be especially effective in producing a number of other neurobiological effects related to the development of addiction-like behavior, including dysregulation of mGluR2/3 receptor function (Allain *et al*., 2017), elevated BDNF levels (Gueye *et al*., 2018), and increased activity in orexin/hypocretin neurons (James *et al*., 2018).

In contrast with the dopaminergic sensitization produced by IntA experience, there are a number of reports that LgA does the opposite – decreases DA function, relative to ShA. For example, following LgA, or other high dose cocaine procedures, the ability of cocaine to inhibit DA uptake, or for electrical stimulation to evoke DA release from the accumbens core, is decreased in tissue slices, as is cocaine-evoked DA overflow measured with microdialysis *in vivo* (Ferris *et al*., 2011; Calipari *et al*., 2013, 2014; Siciliano *et al*., 2016). It may seem surprising, therefore, that in the present study a single, self-administered IV injection of cocaine increased DA to the same extent in rats with LgA or ShA experience – that is, there was no evidence for tolerance. It is not clear what accounts for the discrepancy – e.g., *ex vivo* vs. *in vivo* measures, experimenter-administered IP cocaine challenge vs. self-administered IV injection, measurement technique, or other methodological differences. However, the present results are consistent with one other study on the effects of LgA experience on DA measured with microdialysis *in vivo*. Ahmed (2003) reported that, relative to ShA, LgA did not decrease the DA response in the nucleus accumbens to either experimenter-administered IV injections of cocaine, or cocaine self-administration. Thus, it seems that LgA experience does not consistently decrease DA activity. It should also be noted that effects may vary considerably as a function of how long after the discontinuation of self-administration rats are tested (e.g., Ferrario *et al*., 2005; Siciliano *et al*., 2016). In addition, Willuhn et al. (2014) reported that the magnitude of a phasic DA response seen following a nosepoke that delivered cocaine progressively decreased with increasing LgA experience, as measured with fast scan cyclic voltammetry. However, this phasic DA response peaked approximately 5 sec after a nosepoke, which is too soon to reflect the pharmacological effects of cocaine (Stuber *et al*., 2005; Aragona *et al*., 2008), and therefore, may not be relevant to the studies discussed above.

One hour after the cocaine injection the cue that had been associated with cocaine was presented and we expected to see a conditioned DA response. But the cocaine cue had no effect in any group, on any neurochemical measure. It is not clear why this was the case, because the cue certainly had motivational properties, as indicated by the cue reinstatement test. However, if there were only a very brief (seconds) and relatively small response it may not have been detectable over the 3 min sampling period used here, and other techniques may be required to study the effects of IntA on such conditioned responses.

## Conclusions

It has been suggested that addiction is characterized by a *hypo*dopaminergic, anhedonic state, and compulsive motivation to seek and take cocaine derives from a desire to overcome this DA deficiency (Dackis & Gold, 1985; Koob & Le Moal, 1997, 2001; Blum *et al*., 2015; Volkow *et al*., 2016). Reports that LgA cocaine self-administration experience reduces DA function have been interpreted as support for this view, especially given that LgA was thought to best model changes in brain and behavior that lead to a transition from casual patterns of drug use to the escalated use that characterizes addiction. However, as reviewed above, the evidence that LgA produces a hypodopaminergic state is equivocal, as is evidence it increases motivation for cocaine. In addition, studies using the more recently developed IntA self-administration procedure support a different theory. The IntA procedure was initially developed because it is thought to better model intermittent patterns of cocaine use in humans, especially during the transition to addiction (Zimmer *et al*., 2012; Allain *et al*., 2015). There is now considerable evidence that IntA produces incentive-sensitization, and is more effective than LgA in producing addiction-like behavior (Kawa *et al*., 2016; Allain *et al*., 2017, 2018; Allain & Samaha, 2018; James *et al*., 2018; Kawa & Robinson, 2018). Although the evidence is limited, and more work is required, the available evidence indicates that IntA experience also sensitizes DA function (Calipari *et al*., 2013, 2015), including the ability of cocaine to increase extracellular DA *in vivo,* as reported here.

In conclusion, studies utilizing the IntA procedure are more consistent with the view that the pathological motivation to seek and take cocaine in addiction is due, at least in part, to a hyper-responsive dopaminergic state, consistent with an incentive-sensitization view of addiction (Robinson & Berridge, 1993; Berridge & Robinson, 2016). Of course, a syndrome as complex as addiction is not going to be reducible to changes in a single neurotransmitter system, or even a single psychological process, and it remains to be seen what other neuropsychological functions are altered by IntA experience (e.g., Allain *et al*., 2017; Gueye *et al*., 2018; James *et al*., 2018). Nevertheless, the growing evidence concerning the importance of pharmacokinetic factors in promoting the development of addiction suggests these need to be given greater consideration in preclinical models of addiction (Allain *et al*., 2015).

## ACKNOWLEDGEMENTS

Funding and Disclosure: The authors declare no conflict of interest. This research was supported by grants PO1 DA031656 and T32 DA007281 from NIDA to TER and RO1 EB003320 to RTK.

## DATA ACCESSIBILITY

Raw data generated by these experiments have been stored on a University of Michigan Server. Data and statistical analyses will be made available upon request.

